# A Synthetic ERRα Agonist Induces an Acute Aerobic Exercise Response and Enhances Exercise Capacity

**DOI:** 10.1101/2022.10.05.510974

**Authors:** Cyrielle Billon, Sadichha Sitaula, Subhashis Banerjee, Ryan Welch, Bahaa Elgendy, Lamees Hegazy, Tae Gyu Oh, Melissa Kazantzis, Arindam Chatterjee, John Chrivia, Matthew E. Hayes, Weiyi Xu, Angelica Hamilton, Janice M. Huss, Lilei Zhang, John K Walker, Michael Downes, Ronald M. Evans, Thomas P. Burris

## Abstract

Repetitive physical exercise induces physiological adaptations in skeletal muscle that improves exercise performance and is effective for the prevention and treatment of several diseases. Here we report the identification of a synthetic agonist for the orphan nuclear receptor ERRα (estrogen receptor-related receptor α), SLU-PP-332, that activates an acute aerobic exercise genetic program in skeletal muscle in an ERRα-dependent manner. SLU-PP-332 increases mitochondrial function and cellular respiration consistent with induction of this genetic program. When administered to mice, SLU-PP-332 increased the type IIa oxidative skeletal muscle fibers and enhanced exercise endurance. These data indicate the feasibility of targeting ERRα for development of compounds that act as exercise mimetics that may be effective in treatment of numerous metabolic disorders and to improve muscle function in the aging.

## INTRODUCTION

Lack of physical activity is a substantial contributor to development and progression chronic diseases including obesity, type 2 diabetes, cardiovascular disease, osteoporosis, dementia and cancer^1^. Exercise is an effective treatment for many chronic diseases including obesity and type 2 diabetes^2^, and when exercise is combined with dietary modifications, this treatment can be more effective than currently available pharmacological therapies^3^. Even a single bout of exercise improves whole-body insulin sensitivity for up to 48h after exercise cessation^4^. Furthermore, a single bout of exercise can increase basal energy expenditure beyond the point of exercise termination^5^. Physical exercise is generally classified as either aerobic (endurance-based high frequency repetition with relatively low load) or anaerobic exercise (resistance, strength-based low frequency repetition with relatively high load). The skeletal muscle is one of the primary tissues that adapts to exercise in order to physically and metabolically acclimatize to the increase in utilization. Physical exercise triggers dramatic changes in skeletal muscle gene and protein expression that drive these physiological adaptations that provide for improved muscle function (strength) and endurance can be detected after single bouts of exercise (acute exercise) and repeated bouts of exercise (training)^6^. Both aerobic and anaerobic/resistance exercise are effective in preventing and treating obesity and diabetes, but each induce distinct physiological adaptations within the skeletal muscle. One of the key adaptations of skeletal muscle that occurs in response to aerobic exercise is an increase oxidative capacity of the tissue via elevated mitochondrial respiratory capacity, which allows for more efficient energy production and improved exercise endurance ^7^.

The estrogen receptor-related orphan receptors (ERRα ERRβ and ERRγ) were the first orphan nuclear receptors to be identified^8^. As their moniker indicates, they are homologous to estrogen receptors (ERα and ERβ); however, they do not bind endogenous ER ligands. While ERs require ligand binding to display transcriptional activity, all three ERRs exhibit ligand-independent constitutive transcriptional activation activity^9^. ERRs are highly expressed in tissues with high energy demand such as skeletal muscle, heart, brain, adipose tissue, and liver^8, 10^. A range of target genes whose transcription is activated by ERRs have been identified that includes enzymes and regulatory proteins in energy production pathways involved in fatty acid oxidation, the TCA cycle, mitochondrial biogenesis, and oxidative phosphorylation^11, 12^.

Although the of ERRα null mice are susceptible to heart failure under stress^13^ they can be maintained to investigate ERRα function. A skeletal muscle specific deletion of ERRα yielded mice that displayed reduced mitochondrial biogenesis and impaired repair^14^. A later study using the whole body ERRα-null mice showed that they had decreased muscle mass and decreased exercise endurance that was associated with impaired metabolic transcriptional programs in the skeletal muscle^15^. A genetic gain of function mouse model with ERRγ overexpressed in the skeletal muscle is consistent with these data with the mice displaying increased mitochondrial biogenesis and lipid oxidation^16^. Interestingly, these mice also displayed an increase in oxidative muscle fibers and increased exercise endurance without endurance training^16^. Rangwala et al. reported similar results with overexpression of ERRγ in muscle and additionally, this group also demonstrated that loss of one copy of ERRγ resulted in decreased exercise capacity and mitochondrial function^17^. ERRβ levels are considerably lower than either ERRα or ERRγ in skeletal muscle and thus ERRβ appears have minimal if any relevance in this tissue^18^.

It has been suggested that ERRα is an intractable drug target based on the collapsed putative ligand binding pocket and lack of success of several high throughput screening campaigns as well as the failures of structure based drug design efforts based on homology of ERRα to ERRγ^19^. Based on the observation that skeletal muscle specific ERRα KO mice as well as the ERRα inverse agonist treated mice display decreased exercise tolerance^15^, we sought to identify ERRα agonists that might act as exercise mimetics. Here, we report the identification of an ERRα agonist that effectively induces an acute aerobic exercise genetic program in skeletal muscle in an ERRα-dependent manner. Mitochondrial function and cellular respiration are also enhanced. Importantly, repeated pharmacological activation of ERR provides appears to mimic repeated bouts of aerobic exercise (training) in terms of increased skeletal muscle oxidative capacity and improved exercise endurance.

## RESULTS AND DISCUSSION

### Characterization of SLU-PP-332 as a novel ERRα agonist

It has been suggested that ERRα is an intractable drug target based on the collapsed putative ligand binding pocket and lack of success of several high throughput screen campaigns as well as the failures of structure based drug design efforts based on homology of ERRα to ERRγ^19^. Based on the observation that skeletal muscle specific ERRα KO mice^13^ as well as the ERRα inverse agonist treated mice display decreased exercise tolerance^15^, we sought to identify ERRα agonists that might act as exercise mimetics. Furthermore, the identification of C29 as an ERRα inverse agonist ^20^ suggested that ERRα may not be an intractable drug target. However, as an inverse agonist, C29 acts to block the constitutive transcriptional activation activity of ERRα and it is uncertain if one could design a small molecule that could act as an ERRα agonist enhancing the strong constitutive activity of the receptor.

Using a rational drug design approach (described in the methods) we optimized the ERRβ/γ selective agonist scaffold, GSK4716, for ERRα activity and gained 50-fold ERRα potency. We utilized the X-ray crystal structure of the ligand binding domain (LBD) of ERRγ bound to GSK4716 (Figs. 1a & 1b and Supplementary Fig. 1)(PDBID: 2GPP)^21^ and subsequently modeled GSK4716 bound to the LBD ERRα in order to assess how we might optimize such binding to design an ERRα agonist (Fig. 1c). In contrast to the first report of GSK4716 displaying no ERRα activity, we observed very weak agonist activity in an ERR cotransfection assay (Supplementary Fig. 2a) and believed that the GSK4716 scaffold may be useful as an initiation point to develop high affinity ERRα agonists. The X-ray structure of ERRγ LBD bound to GSK4716 is the only X-ray crystal structure for any ERR bound with an agonist ligand (Fig. 1b)^21^.

**Figure 1.**
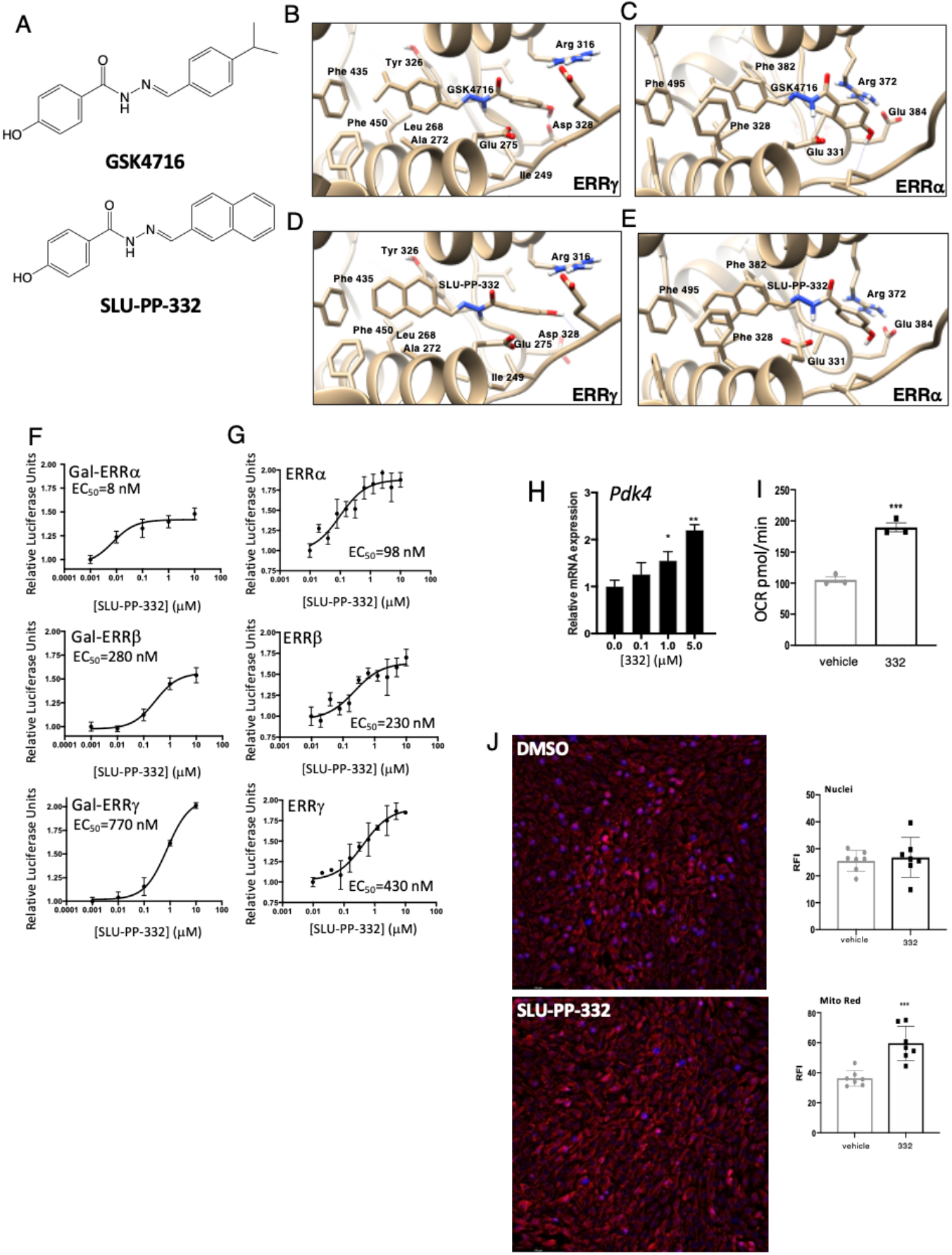
SLU-PP-332 is a synthetic ERRα selective agonist. **(A)** Chemical structures of GSK4716 (top) and SLU-PP-332 (bottom). **(B)** Schematic illustrating the X-ray co-crystal structure of GSK4716 bound to the ERRγ LBD (PDBID: 2GPP). Models of ERRα bound to GSK4716 **(C)**, ERRγ LBD bound to GSK4716 **(D)** and ERRα bound to SLU-PP-332 **(E)**. Results of ERRα, ERRβ or ERRγ co-transfection assays Gal4-DBD ERR LBD chimeric receptors **(F)** and full length ERRs **(G)** in HEK293 cells illustrating the activity of SLU-PP-332. **(H)** Effects of SLU-PP-332 dose-response treatment (24h) on *Pdk4*, in C2C12 cells, n=3. **(I)** Maximal mitochondrial respiration values analyzed by Seahorse of C2C12 cells treated with DMSO (gray bar) or SLU-PP-332 (black bar), n=3. **(J)** Mitotracker red staining of C2C12 cells under proliferative conditions treated with DMSO or SLU-PP-332 (10 µM) for 24h. Bar graph represents nuclei intensity (left) and mitotracker red staining (right). p<0.05, ** p<0.01, $ p<0.05.

In this structure, the agonist GSK4716 binds in a previously unidentified pocket dubbed the “agonist” pocket near the solvent exposed surface of the receptor (Supplementary Fig. 1)^21^. The phenolic hydroxyl group of GSK4716 interacts with Asp328 (Fig. 1b) while the carbonyl of the acyl hydrazone bridges to two water molecules, one of which interacts with Arg316 and the other water molecule interacts with Leu309 (Supplementary Fig. 1). Molecular modeling of GSK4716 in the LBD of ERRα followed by energy minimization, to refine the protein-ligand complexes, reveals similar interactions to that observed with ERRγ X-ray crystal structure (Fig. 1c). Of particular importance, a phenylalanine residue, Phe328, is present in the LBD of ERRα which corresponds to Ala272 in ERRγ and Ala274 in ERRβ. Our strategy to design high affinity ERRα agonists was based on optimizing interaction ligand interactions with Phe328 that is specifically in ERRα. We employed a strategy to optimize GSK4716 based on converting the isopropyl phenyl group of GSK4716 to a more hydrophobic moiety that could potentially gain affinity by interacting with the Phe328 in ERRα. Molecular modeling of a compound with a naphthalene substituent in place of the isopropyl phenyl group (SLU-PP-332; Fig. 1a bottom and Fig. 1c and Supplementary Fig. 3) in the LBD of ERRγ and ERRα predicted the newly added phenyl group to make π-π interactions with Phe435 (ERRγ) or Phe495 and Phe328 (ERRα). We hypothesized that this modification would improve the affinity of the ligand in both receptors, but particularly towards ERRα due to a potential π-π stacking interaction between the ligand naphthalene group and Phe328 (the corresponding alanine residue in ERRβ and ERRγ is unable to make such interactions). As predicted, SLU-PP-332 gained substantial ERRα potency (Figs 1d & 1e) in addition to a moderate increase in potency for ERRβ and γ (Figs. 1f & 1g). SLU-PP-332 displayed considerable enhanced potency over GSK4716 for all three ERRs but was particularly more potent activating ERRα (Figs. 1f & 1g). SLU-PP-332 was nearly 100-fold more potent at ERRα than ERRγ in the chimeric Gal4 DBD ERR LBD – luciferase reporter cotransfection assay in HEK293T cells (ERRα EC_50_=8nM, ERRβ=280nM, ERRγ=770nM)(Fig. 1f). In a cell-based contransfection assay utilizing full length ERRs and a ERR response element enhancer driven luciferase reporter SLU-PP-332 was still ERRα selective displaying 4.4-fold selectivity for ERRα over ERRγ (ERRα EC_50_=98nM, ERRβ=230nM, ERRγ=430nM) (Fig. 1g). SLU-PP-332 was selective for the ERRs as it did not alter the activity of either ERα or ERβ, or other nuclear receptors in co-transfection assays (Supplementary Fig. 2b). Direct binding of SLU-PP-332 to the ERRα LBD was assessed by limited proteolysis where the LBD is subjected to chymotrypsin in the presence and absence of SLU-PP-332 and the ability of the drug to “protect” fragments of the LBD from digestion due to a conformational change in the LBD is detected^22^. As shown in Supplementary Fig. 2c, SLU-PP-332 dose-dependently protects a fragment of the ERRα LBD from protease digestion consistent with direct binding of the drug to ERRα. Direct binding of SLU-PP-332 to ERRγ was also confirmed using differential scanning fluorimetry, where the compound dose-dependently increased the thermal stability of the purified ERRγ LBD (Supplementary Fig. 2d)

### SLU-PP-332 increases the expression of an ERR target gene and enhances mitochondrial respiration in C2C12 myocytes

We next examined whether SLU-PP-332 could increase the expression of an ERR target gene in the C2C12 myoblast cell line. We noted a clear increase in the expression of a well-characterized ERR target gene, *pyruvate dehydrogenase kinase 4* (*Pdk4*)^23^ with SLU-PP-332 treatment (Fig. 1h). Overexpression of ERRγ in C2C12 myocytes has been demonstrated to enhance mitochondrial respiration and pharmacological inhibition of ERRα suppresses mitochondrial respiration in these cells^24^, thus we hypothesized that SLU-PP-332 would enhance mitochondrial respiration. Proliferating C2C12 cells treated with SLU-PP-332 for 24 hours exhibited an increase in maximum mitochondrial respiration relative to cells treated with vehicle (Fig. 1i and Supplementary Fig. 4). Furthermore, we observed that SLU-PP-332 treatment substantially induced mitochondrial biogenesis in proliferating C2C12 cells based on staining with mitotraker red (Fig. 1j).

### SLU-PP-332 induces enhances exercise endurance in mice

We sought to determine if SLU-PP-332 could potentially be used as a chemical probe to evaluation activation of ERR function *in vivo* thus we first assessed in vivo exposure after intraperitoneally (i.p.) administration in mice (Fig. 2a). Mice were administered SLU-PP-332 (30mg/kg, i.p.) and plasma and muscle were collected 2-and 6-hours post administration and analyzed by mass spectrometry. Two hours after administration levels of SLU-PP-332 were highest in skeletal muscle (∼0.8 µM) while levels in the plasma were lower (∼0.1 µM) (Fig. 2a). We observed no overt toxicity in mice administered SLU-PP-332 (50 mg/kg b.i.d., i.p.) for 10 days, which is consistent with the normal complete blood count and electrolyte levels^25^ (Supplementary Table 1). We also observed no significant alterations in serum creatine kinase suggesting the lack of skeletal muscle toxicity (Supplementary Fig. 5)

**Figure 2.**
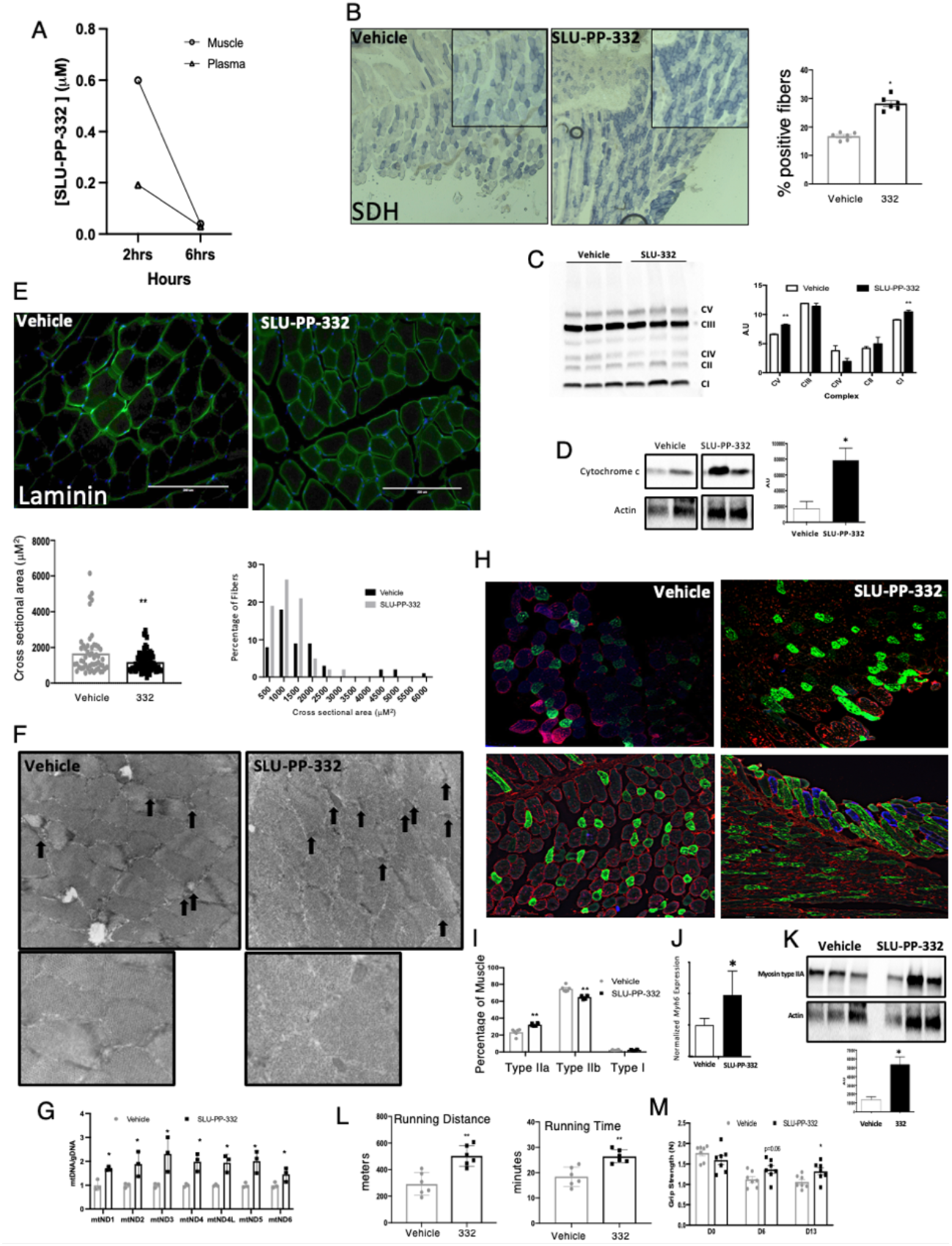
SLU-PP-332 increases oxidative fibers in skeletal muscle and improves exercise endurance. **(A)** Pharmacokinetic analysis of SLU-PP-332 displaying muscle and plasma levels of the compound at 2 and 6 h post 30mg/kg, i.p. in 10/10/80 DMSO, Tween, PBS. n=3. **(B)** Immunochemical analysis of succinate dehydrogenase (SDH) from quadriceps of mice administered vehicle (white bars, n=8) or SLU-PP-332 50mg/kg, b.i.d (black bar, n=8). The bar graph represents the quantification of SDH positive muscle fibers. OXPHOS complex **(C)** and Cytochrome C **(D)** protein levels from quadriceps from mice dosed with vehicle (white bars, n=7) or SLU-PP-332 50mg/kg, b.i.d (black bar, n=7). The bar graph represents the quantification of expression. Immunochemical analysis of laminin **(E)**, Electron microscopy of quadriceps **(F)** (Black arrows illustrated identified mitochondria). **(G)** Analysis of mitochondrial DNA levels (relative to nuclear DNA) from quadriceps. Immunochemical analysis of muscle fiber types **(H)** stained sections (n=6 per group) from quadriceps of mice administered vehicle (white bars, n=8) or SLU-PP-332 50mg/kg, b.i.d (black bar, n=8). Myosin IIa is green, myosin IIb is red and myosin I is blue. The bar graph represents the quantification of fiber cross sectional area (e lower panel) and percentage of fiber types **(I)**. Myosin heavey chain gene **(J)** and protein **(K)** expression from quadriceps of mice treated with vehicle or SLU-PP-332 50mg/kg, b.i.d (n=6 for gene expression and n=3 for protein). The bar graph represents the quantification of expression. Mitochondrial DNA content (g) from the same mice (n=3 per group). **(L)** Running distance (left panel) and running time (right panel) of mice treated with an acute dose of vehicle (gray bar) or SLU-PP-332 (50mg/kg, black bar) for 1h before running (n=6 per group). **(M)** Grip strength test from mice treated with vehicle (gray bar) or SLU-PP-332 (50mg/kg, black bar), before dosing (D0), after 6 days of dosing (D6) or after 13 days (D13) (n=8). p<0.05, ** p<0.01, *** p<0.001, **** p<0.0001.

We next examined the effects of chronic SLU-PP-332 treatment on muscle physiology and function. Three-month old C57BL/6J mice were administered SLU-PP-332 for 15 days (30 mg/kg, b.i.d., i.p.) followed by examination of quadricep muscle histology. To avoid the effect of ERR on facultative thermogenesis and cold tolerance^26^ we maintained the mice at thermoneutrality (30°C). Histology was performed on unfixed muscle and stained for hematoxylin and eosin (Supplementary Fig. 6a) succinate dehydrogenase (SDH) activity (Fig. 2b). Mice treated with SLU-PP-332 displayed a more oxidative muscle phenotype (greater SDH staining) (Fig. 2b). We also assessed expression of key proteins within the oxidative phosphorylation complexes and observed an increase in Complex I (NDUFB8) and Complex V (ATP5A) expression in the gastrocnemius muscle in response to SLU-PP-332 treatment (Fig. 2c). Consistent with these observations we also observed an increase in cytochrome c protein expression in response to SLU-PP-332 treatment in the gastrocnemius muscle (Fig. 2d). Sections were also stained for laminin and consistent with an increased oxidative phenotype as the myofibers were smaller in diameter^27^ in SLU-PP-332 treated mice (Fig. 2e). Electron micrographs of the quadriceps display an increase in mitochondria content in muscles from drug treated mice compared to vehicle-treated (Fig. 2f). Mitochondrial DNA concentrations in the skeletal muscle were also increased (relative to nuclear DNA) consistent with an increase in mitochondrial number (Fig. 2g). We also noted that SLU-PP-332 treatment resulted in increased oxidative type IIa muscle fibers (Fig. 2h & 2i). This observation aligned with an increase in expression of *Myh6*, which encodes a myosin heavy chain subtype that is associated with type IIa muscle fibers^28^(Fig. 2j). These data indicate that pharmacological activation of ERRs leads to increased oxidative capacity of skeletal muscle and an increase in type IIa muscle fibers and suggested that treatment of mice with SLU-PP-332 may lead to an increase in exercise endurance. In order to assess this, we treated sedentary mice with SLU-PP-332 or vehicle for 7 days (b.i.d., i.p. 30mg/kg) and subjected them to exercise until exhaustion on a rodent treadmill. Plasma glucose levels were monitored following the exercise to confirm exhaustion (Supplementary Fig. 6b). As shown in Fig. 2l, mice treated with the ERR agonist were able to run ∼70% longer and ∼45% further than vehicle treated mice. In a separate study where sedentary mice were treated in an identical manner except longer duration (two-weeks) we noted an increase in grip strength as well (Fig. 2m).

We subsequently assessed gene expression from gastrocnemius and quadricep muscle from mice treated with SLU-PP-332 or vehicle (30 mg/kg, b.i.d., i.p., 10 days) to assess alterations in gene expression due to drug treatment. Muscles were obtained 3h post the final administration and global changes in gene expression were assessed by RNA-Seq. Treatment of sedentary mice with SLU-PP-332 induced an array of genes that substantially overlapped with genes previously shown to be upregulated in response to acute aerobic exercise^29^ (Fig. 3a). This was brought to our attention immediately due to the gene *Ddit4/Redd1* gene (DNA Damage Inducible Transcript 4/Regulated in development and DNA damage responses 1) as the most upregulated gene in both the gastrocnemius and quadricep muscles (Fig. 3a). *Ddit4* expression has been demonstrated to increase transiently following acute aerobic exercise^30 31^ and is responsible for directing an acute aerobic exercise-mediated gene expression signature^32^. Importantly, DDIT4 is critical for exercise adaptation and skeletal muscle mitochondrial respiration as *Ddit4*^*-/-*^ mice display reduced mitochondrial respiration in skeletal muscle and impaired exercise capacity^33^ as well as substantially impaired glucose tolerance^34^. Expression of *Ddit4* is also suppressed in skeletal muscle of rhesus monkeys that are obese and exhibit symptoms of metabolic syndrome^35^. We also noted an increase in the expression of a key gene induced as a component of the *Ddit4*-dependent acute aerobic exercise program, *Slc25a25*^32^. SLC25A25 is an ATP-Mg^2+^/P_i_ inner mitochondrial membrane solute transporter and plays a role in loading nascent mitochondria with nucleotides. Mice deficient in *Slc25a25* expression display reduced metabolic efficiency and decreased exercise endurance^36^. Reduced *Slc25a25* expression leads to lower mitochondrial respiration^36^ while enhanced Slc25a25 has been associated with increased mitochondrial respiration^37^. Of the ∼20 genes that have been reported to be the highest up-regulated in response to acute aerobic exercise in the gastrocnemius^29^, 44% of these are included in the significantly upregulated genes from the gastrocnemius in response to SLU-PP-332 treatment (Fig. 3b & 3c). Importantly, *Ddit4* and *Slc25a25* are included within this series of genes that are both upregulated by acute aerobic exercise and SLU-PP-332 treatment (Fig. 3b). Using the significant gene sets from gastrocnemius and quadriceps muscles (FDR<0.05, FC>2), we analyzed the potentially associated transcription factors in our treated samples using the EnrichR tool. Notably, we observed an ESRRA (ERRα) associated gene set in the data analyzed from both gastrocnemius and quadriceps muscle (Supplementary Fig. 7). This analysis suggests that SLU-PP-332 treatment is tightly associated with ERRα-dependent genes. Furthermore, we examined whether ERRα directly binds to overlapped genes from gastrocnemius and quadriceps muscle samples by examining previously published ChIP-seq data that utilized C2C12 myocytes^38^. We found ERRα is recruited near or within these genes that are included within the acute aerobic exercise genetic program including *Ddit4, Slc25a25, Mypn, Nr4a1, Ide, Hbb-bt, Hba-a1, Gadd45g* and *Tsc22d3* (Supplementary Fig. 8). Pathway analysis of genes upregulated by SLU-PP-332 treatment demonstrated considerable overlap in the gastrocnemius and quadricep muscles. Five of the of the top 10 pathways (Wikipathways 2019) significantly upregulated were shared between the muscles types and are shown in Fig. 3d. Interestingly, the exercise-induced circadian regulation pathway was affected in both muscles with *Per1* and *Per2* both significantly upregulated (Fig. 3e & 3f). The expression of *Per1* and *Per2* has been demonstrated to be induced by acute aerobic exercise previously and, most importantly, the induction of *Per1* is completely dependent on DDIT4^39^. Although SLU-PP-332 treatment modulated *Per1* expression in skeletal muscle, we observed no alterations in circadian locomotor activity with drug treatment (Supplementary Fig. 9b). *Foxo1* is also a well-characterized gene induced by acute endurance exercise in skeletal muscle^40^ and was also induced by SLU-PP-332 treatment (Fig. 3e & 3f). δ-aminolevulinate synthase 2 (*Alas2*) was also upregulated by SLU-PP-332 treatment and ALAS activity in skeletal muscle has been previously demonstrated to be enhanced by exercise^41^. ALAS catalyzes the rate limiting step in heme synthesis and there are two forms of the enzyme encoded by the *Alas1* and *Alas2* genes. *Alas1* is ubiquitously expressed is its expression in skeletal muscle is induced by exercise^42^ whereas *Alas2* expression is generally characterized as erythroid cell specific. However, *Alas2* has been demonstrated to be expressed in skeletal muscle and its expression is modulated by macronutrients in the diet^35^. With this data in hand indicating that SLU-PP-332 treatment enhanced exercise endurance and induced a gene program similar to acute aerobic exercise in skeletal muscle, we sought to compare the effects of acute SLU-PP-332 treatment to acute aerobic exercise in terms of induction of *Ddit4* and *Slc25a25*. Three-month old C57BL/6J male mice were administered a single dose of SLU-PP-332 (30 mg/kg; i.p.) or vehicle and compared to mice that were sedentary or subject to acute aerobic exercise (run for 40 minutes on a rodent treadmill). *Ddit4* and *Slc25a25* gene expression from the gastrocnemius muscle was assessed at 1, 3 and 6h post initiation of the run or drug administration (from distinct groups of mice). At 1h post run or treatment initiation, *Ddit4* expression was ∼6-fold higher in the exercised group and, most importantly, we noted that SLU-PP-332 treatment of sedentary mice induced an ∼3-fold increase in this gene (Fig. 3g top). Mice that received both the drug and were exercised displayed an additive effect on *Ddit4* expression (∼11-fold increase; Fig. 3g top). All of these effects were transient, and the effects were not observed in the later time points (Fig. 3g top). Acute aerobic exercise and SLU-PP-332 treatment led to equivalent 11-fold increases in *Slc25a25* expression after 1 h and like *Ddit4*, the effect was lost after 3 h (Fig. 3g bottom). Thus, induction of these two acute aerobic exercise program genes by either SLU-PP-332 or exercise was similar in terms of magnitude and duration. We also examined DDIT4 protein expression from the quadricep muscle 1 and 3h post SLU-PP-332 treatment and noted a time-dependent increase in expression reaching a level 3.5X higher after 3h (Fig. 3c). We also examined ERRα expression at 1h and 3h post administration of the drug and observed no change in expression indicating that the kinetics of the response was not due to changes in ERRα expression (Supplementary Fig. 9a)

**Figure 3.**
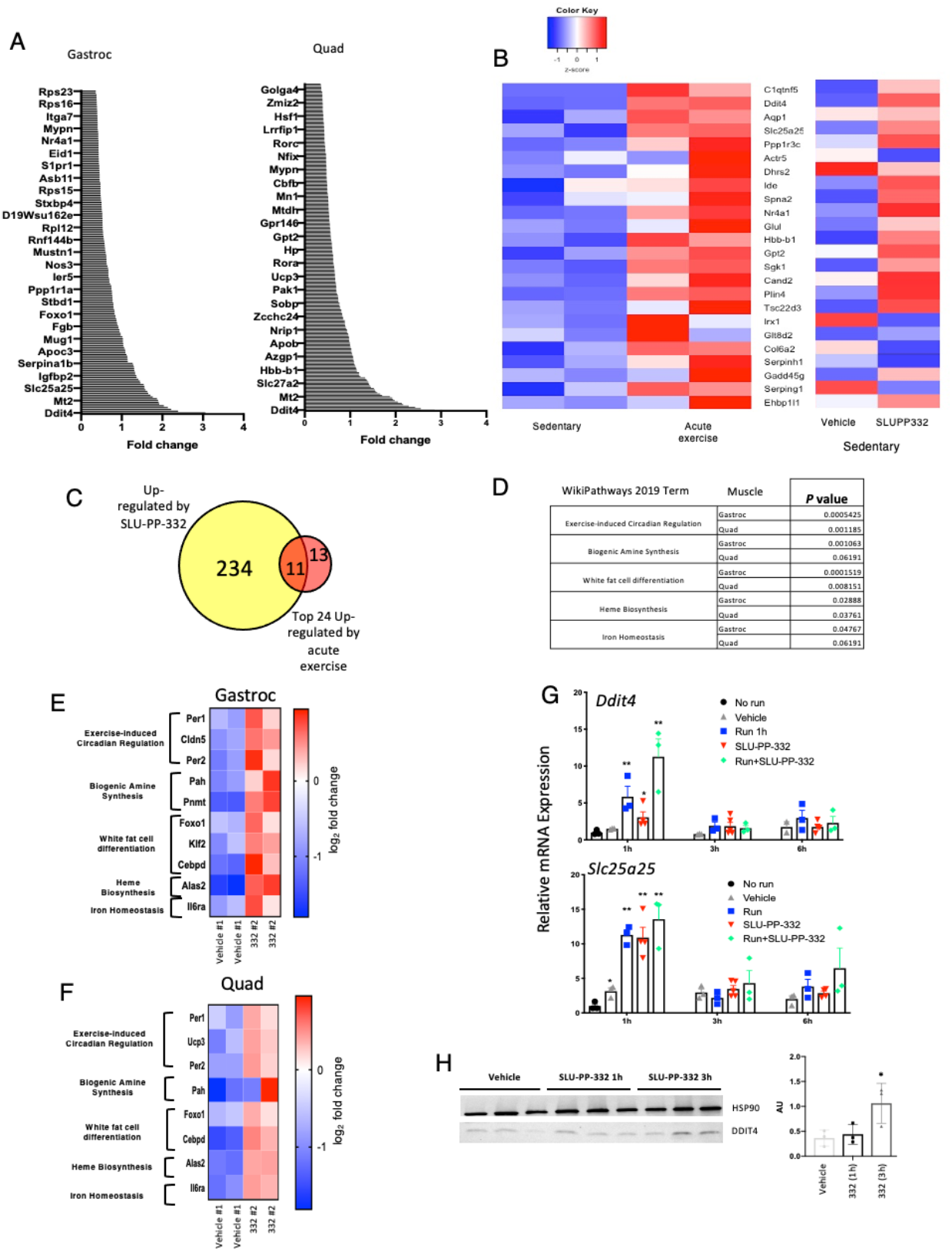
SLU-PP-332 induces acute aerobic exercise genetic program in skeletal muscles. Results from RNA-seq studies from the gastrocnemius of mice treated with vehicle or SLU-PP-332. **(A)** Ranking of the top 25 genes up-regulated by SLU-PP-332 in both gastrocnemius and quadriceps from mice treated with SLU-PP-332 (50mg/kg, b.i.p) for 10 days. Heat map **(B)** and Venn Diagram representation **(C)** of an analysis of results from Sako et al^29^ comparing sedentary mice (b, left panel) to acute exercised mice and vehicle to SLU-PP-332 treated mice (b, right panel). Pathway analysis (**D)** of genes induced from quadricep and gastrocnemius muscles after treatment with SLU-PP-332 in mice. **(E)** Heatmap of genes from pathways identified in (D) above. **(g)** *Ddit4* (upper panel) and *Slc25c25* expression (lower panel) from quadriceps from mice treated with vehicle (gray triangle), SLU-PP-332 (50mg/kg, IP, red triangle), SLU-PP-332 in combination of 45 min of running (green circle) or no treatment (black circle). Mice were euthanized as indicated 1h, 3h or 6h after treatment (n=6 per group). (h) Ddit4 protein expression from quadriceps from mice treated with SLU-PP-332 (50mg/kg, IP). Mice were euthanized as indicated 1h or 3h (n=3 per group). The bar graph represents the quantification of expression. p<0.05, ** p<0.01, *** p<0.001, **** p<0.0001.

### *Ddit4*, a regulator of an acute aerobic exercise genetic program, is an ERRα specific target gene

We next sought to characterize ERR regulation of *Ddit4* in greater detail using the C2C12 myoblast cell line. Using both a QPCR array (Fig. 4a) and direct QPCR (Fig. 4b), we found that *Ddit4* gene expression was induced in the C2C12 myoblast cell line by SLU-PP-332 treatment and the induction was detected in as little as 1h (Fig. 4a & 4b). Analysis of previously published ERRα ChIP-seq data from C2C12 cells^38^ revealed ERRα occupancy in the 5’ region and intragenic regions of the *Ddit4* gene (Fig. 4c) as well as within many other of the other genes regulated by SLU-PP-332 including *Slc25a25, Mypn, Nr4a1, Ide, Hbb-bt, Hba-a1, Gadd45g* and *Tsc22d3* (Supplementary Fig. 7). Multiple putative ERREs were identified the promoter and intragenic regions of *Ddit4* (Fig. 4d). We assessed the promoter region bound by ERRα identified in the ChIP-seq data containing a putative ERRE that conferred SLU-PP-332 responsiveness to a luciferase reporter gene when cotransfected into HEK293 cells along with ERRα, consistent with *Ddit4* functioning as a direct ERRα target gene (Fig. 4e) We also observed that *Ddit4* expression was induced in primary mouse myocytes (derived from quadriceps) by acute SLU-PP-332 treatment (2h) (Fig. 4f) but the effect was not observed after 24h treatment (Fig. 4g) reminiscent of the transitory effect we observed in vivo and consistent with the transient induction of genes in the acute aerobic exercise genetic program. The effect of SLU-PP-332 on *Ddit4* expression was dependent on ERRα since the effect was lost in myocytes derived from ERRα or ERRα/γ null mice but was retained in ERRγ null myocytes (Fig. 4f). ERRβ is not expressed in these cells (Supplementary Fig. 9c). SLU-PP-332 induced expression of *Slc25a25* in a pattern identical to *Ddit4* (Fig. 4h). *Slc25a25* responsiveness was completely ERRα-dependent and was transient with an effect noted at 2h but not at 24h post treatment (Fig. 4i). These results in the primary myocytes suggest that the effects of SLU-PP-332 on induction of the acute aerobic exercise genes such as *Ddit4* are mediated via ERRα and not ERRβ or ERRγ. In order to investigate this in the context of the whole animal, we treated mice with a muscle specific KO of ERRα with SLU-PP-332. mERRα^fl/fl^ or mERRα^-/-14^ were treated for 14 days with SLU-PP-332 (b.i.d., i.p, 25mg/kg) and then subjected to exercise until exhaustion. mERRα^fl/fl^ treated with SLU-PP-332 exhibited significantly enhanced exercise endurance while the mERRα^-/-^ treated with SLU-PP-332 displayed exercise endurance equivalent to vehicle treated mERRα^-/-^ mice (Fig. 4j). Moreover up-regulation of *Ddit4* in quadriceps was observed only in mERRα^fl/fl^ treated with SLU-PP-332 but not in the mERRα^-/-^ treated group (Fig. 4k). Additionally, two other genes that we identified as upregulated in response to SLU-PP-332 treatment in skeletal muscle, *Per1* and *Alas2* (Fig.3f), were significantly induced in the mERRα^fl/fl^ mice by SLU-PP-332 treatment but not in the mERRα^-/-^ mice (Fig. 4l & 4m). These data illustrating the ability of administration of SLU-PP-332, a compound that induces an acute exercise genetic program via activation of ERRα, to increase exercise endurance are consistent with studies demonstrating that both *Ddit4* and *Slc25a25* are key regulators of mitochondrial function and mice with null mutations in either of these genes exhibit substantially reduced exercise endurance^33, 36, 37^.

**Figure 4.**
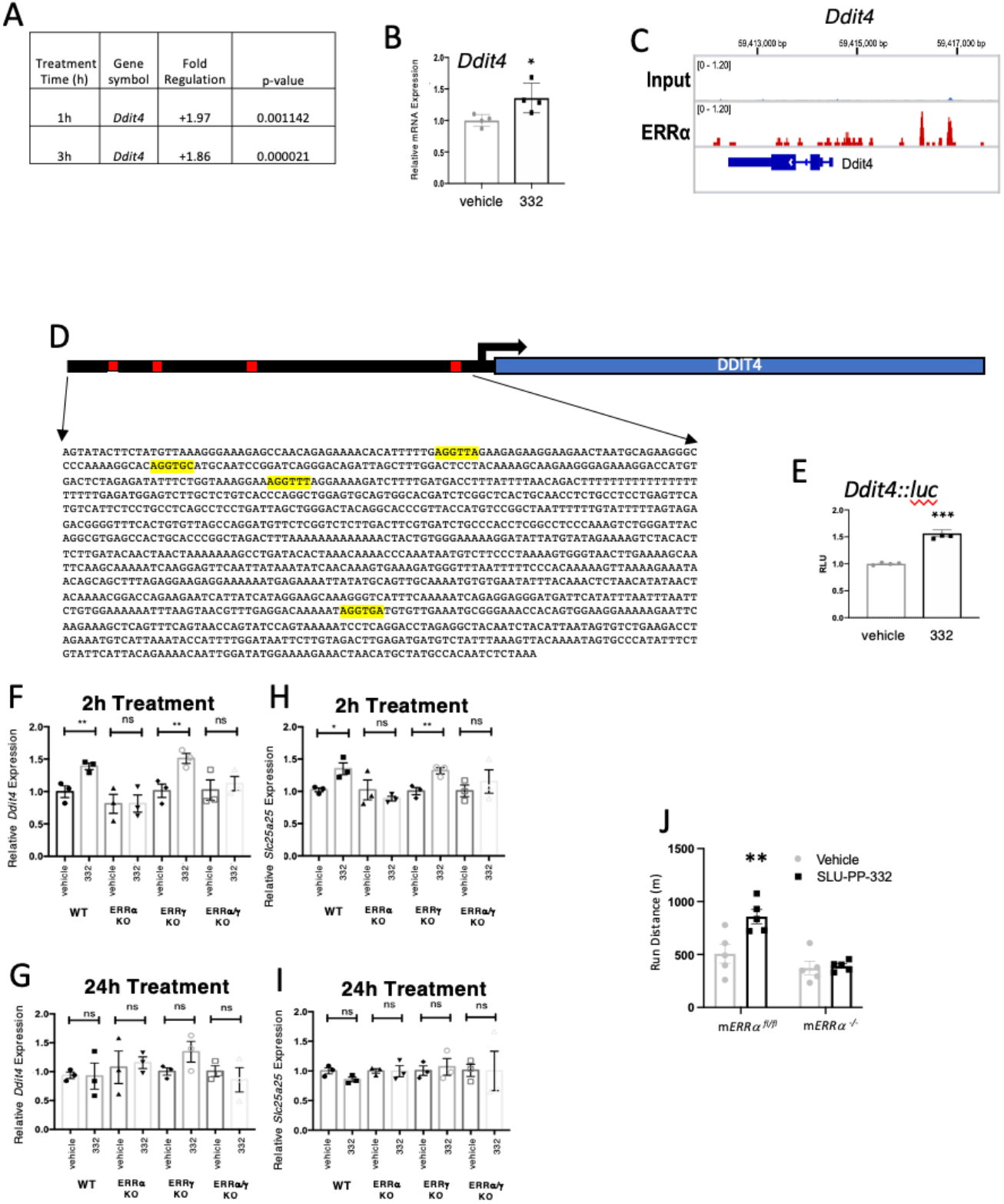
SLU-PP-332 induces *Ddit4* expression and enhances exercise endurance in an ERRα-dependent manner. **(A)** Effects of SLU-PP-332 treatment (10 µM) for either 1 or 3 h on *Ddit4* expression in C2C12 cells detected using the Qiagen RT^2^ PCR array **(B)** Effects of SLU-PP-332 (10μM) treatment of C2C12 for 3h gene on *Ddit4* expression (n=3). **(C)** ERRα binding locations near and within the *Ddit4* gene identified in by ChIP-seq in C2C12 cells. **(D)** Schematic representation of luciferase reporter containing the putative ERRα binding site from *Ddit4* identified in (c). **(E)** Cotransfection assay in HEK293T cells with full length ERRα (including SLU-PP-332 (10µM)). *Ddit4 and* Slc25a25 expression are transiently induced in primary myocytes by SLU-PP-332 in an ERRα-dependent manner. *Ddit4* expression in primary ERR WT, ERRα KO, ERRγ KO and ERRα/γ KO myoblasts treated with DMSO or SLU-PP-332 (1 µM) for 2hrs **(F)** or 24hrs **(G)**. *Slc25a25* expression in primary ERR WT, ERRα KO, ERRγ KO and ERRα/γ KO myoblasts treated with DMSO or SLU-PP-332 (1 µM) for 2hrs **(H)** or 24hrs **(I). (J)** Running endurance of mERRα^fl/fl^ vs. mERRα^-/-^ mice doses with vehicle or SLU-PP-332. p<0.05, ** p<0.01, *** p<0.001, **** p<0.0001

## CONCLUSIONS

The ERRs play important roles in regulation of energy metabolism and fuel selection. Loss of ERRα or ERRγ function leads to reduced muscle oxidative function and reduced functional endurance^15, 43^, thus pharmacological activation of these receptors may lead to beneficial metabolic effects associated with increased skeletal muscle activity for treatment of metabolic diseases. In this study, we characterize the ability of an ERRα synthetic agonist with ∼4-5-fold selectivity over ERRγ (SLU-PP-332) to function as an exercise mimetic and improve muscle and metabolic function both *in vitro* and *in vivo*. SLU-PP-332 treatment induces the expression of DDIT4 via specific activation of ERRα. DDIT4 is a key protein that is induced after short bouts of aerobic exercise that is responsible for inducing an acute aerobic exercise genetic program that leads to a range of physiological adaptations to exercise^32^. We found that *Ddit4* is a direct ERRα target gene and previous data indicating that *Ddit4* null mice display reduced exercise endurance^33^ is consistent with our results indicating that SLU-PP-332 treatment, which induces *Ddit4* expression, enhances exercise endurance. The array of genes that are induced by both SLU-PP-332 and acute aerobic exercise have been linked mechanistically to improved exercise endurance, increased fatty acid oxidation and/or improved metabolic efficiency that are all physiological components of the adaptive response to exercise. Of course, the key gene we examined, *Ddit4*, is associated with mitochondrial function and improved exercise endurance^33^ and another gene within this program we examined, *Slc25a25* is also similarly associated with these endpoints^36^. Most importantly, the effects of SLU-PP-332 on exercise endurance is completely dependent on ERRα as mice with muscle specific deletion of this receptor are refractory to the improved performance.

Previously, it had been reported that skeletal muscle overexpression of ERRγ in mice led to improved endurance^16^; however, we noted substantial differences in the array of genes regulated by overexpression of ERRs compared to pharmacological activation of ERRs. The acute aerobic exercise gene program was not identified when either ERRα or ERRγ were overexpressed or knocked-out^9, 11^. We believe that transient activation of ERRs may be quite distinct than either chronic overexpression or complete loss of the receptor. Furthermore, overexpression of ERR(s) likely provides for an unremitting level of elevated transcriptional activity that cannot be mimicked by pharmacological activation of this receptor class that already displays strong constitutive transactivation activity. Multiple bouts of aerobic exercise (2h/day for 8 days on rodent treadmill) has been shown to induce ERRα (∼1.5-fold) and ERRγ (∼2.1-fold) expression within the gastrocnemius muscle in mice^17^. Short-term aerobic exercise (cycling) in humans has been shown to induce ERRα (mRNA and protein expression) in skeletal muscle (m. vastus lateralis) to a similar extent (1.5 - 2 fold)^6^. These data suggest that the ∼2-fold increase in the ERR transcriptional activity that we observe with SLU-PP-332 treatment is likely more similar to physiological changes in ERR activity induced by exercise than experimental models of skeletal muscle overexpression of ERR(s) or VP-16 ERR fusion proteins. Thus, pharmacological activation of ERR may be more closely aligned with driving physiological changes that are similar to normal exercise adaptation such as induction of the acute aerobic exercise response rather than chronic overexpression of ERR or similar key regulator proteins.

In summary, activation of ERRα by SLU-PP-332 as an exercise mimetic inducing an acute aerobic exercise program that leads to an array of physiological adaptations that are associated with exercise including increased oxidative fibers in muscle, increased fatty acid oxidation and enhanced exercise endurance. Several nuclear receptors such as LXR, FXR, PPAR α, PPARδ, PPARγ, and REV-ERB, among others, have been evaluated or utilized as targets for compounds for treatment of metabolic disease. However, only pharmacological activation of REV-ERB and PPARδ have been reported to have exercise mimetic activity^44^. Interestingly, activation of the acute aerobic exercise program appears to be unique for ERR agonists as this was not reported for PPARδ or REV-ERB. ERRα targeted compounds that increase the metabolic performance of skeletal muscle may hold utility in the treatment of metabolic diseases and diseases of muscle atrophy and dysfunction including muscular dystrophy and sarcopenia.

## METHODS

### Molecular Modeling

All four models of ERRγ and ERRα bound with GSK4716 or SLUPP332 were built from the X-ray crystal structure of ERRγ-GSK4716 (PDB:2gpp)^21^. SLUPP332 was modelled by modifying the isopropyl group into a naphthalene group using Maestro (Schrodinger Release 2019-1: Maestro, Schrodinger, LLC. New York, NY). The initial X-ray structure has two water molecules bridging the ligand and the protein residues and they were kept in the ligand binding pocket in each model. Each system was first energy minimized using the steepest descent and conjugate gradient methods with keeping the ligand and the bridged water molecules constrained. The constraints were removed and then each system was energy minimized entirely in Amber^45^. Tleap module was used to neutralize and solvate the complexes using an octahedral water box of TIP3P water molecules. The FF14SB forcefield parameters were used for all receptor residues and the general amber force field was applied to ligand residues^46^. Non-bonded interactions were cut off at 10.0 Å, and long-range electrostatic interactions were computed using the particle mesh Ewald (PME). Ligands were modelled using Maestro and pictures were generated using UCSF Chimera and Maestro^47^. After energy minimization, the water molecules left the pocket and the acyl hydrazine made alternative hydrogen bonding interaction with the protein back bone residues. The phenolic hydroxyl in the ERRγ-GSK4716 model maintained similar hydrogen bonding interaction with Asp328 as the starting X-ray structure. Energy minimization using MacroModel and the OPLS3 force field yielded similar results (Schrodinger Release 2019-3: MacroModel, Schrodinger, LLC. New York, NY) ^48^. Although, the phenolic hydroxyl group in the other three models made hydrogen bonding interactions with different protein residues, it remained in similar position as in the ERRγ-GSK4716 X-ray structure near the solvent exposed surface of the protein (Figure 1). We used the MM/GBSA^49^ method to estimate the binding free energies of GSK4716 and SLU-PP-332 to both receptors (Supplementary Table 1). MM/GBSA, an end point energy calculation method used for estimating relative binding free energies, is particularly useful for ligand ranking and optimization in the process of drug discovery^50^. The binding of GSK4716 and SLU-PP-332 to ERRα and ERRγ is enthalpy driven with a negative total binding free energy indicating favorable binding (Supplementary Table 2). ΔH corresponds to the favorable affinity contribution, while ΔS is the entropy and reflects the decrease in conformational freedom in the protein ligand complex. In the case of ERRγ the enthalpy contribution of GSK4716 to the total binding free energy is more favorable than SLU-PP-332, however the entropy penalty is greater in the case of GSK4716 over SLU-PP-332 resulting in similar total binding free energies with a difference of 1.5 kcal/mol in favor of SLU-PP-332 (Supplementary Table 1). However, in the case of ERRα, the enthalpy contribution of SLU-PP-332 is more favorable and the entropy penalty less resulting in a more favorable total binding free energy (6.6 kcal/mol difference between SLU-PP-332 and GSK4716) (Supplementary Table 2). Based on these calculations, SLU-PP-332 was predicted to have higher affinity for both receptors and particularly towards ERRα, with the reduction of the unfavorable entropic contribution associated with ligand binding the main contributor towards the improved affinity. Additionally, several analogues of GSK4716 where the isopropyl phenyl group was replaced were tested and the naphthalene substituent (SLU-PP-332) was considerably more potent than any others (Supplementary Table 3).

### Synthesis and preparation of SLU-PP-332

(E)-4-hydroxy-N’-(naphthalen-2-ylmethylene)benzohydrazide – To a solution of 2-naphthaldehyde (1.0 g, 6.6 mmol) in toluene (100 mL) was added 4-hydroxybenzohydrazide (1.1 g, 6.6 mmol) portion wise. The mixture was allowed to stir for 18h at reflux. Solid precipitated, which was recrystallized from a 1:9 mixture of methanol and ether to obtain the title compound as white solid (1.3 g, 68%); 1H NMR (400 MHz, DMSO-d6) δ 11.81 (s, 1H), 10.18 (s, 1H), 8.63 (s, 1H), 8.12 (s, 1H), 8.06 – 7.84 (m, 6H), 7.54 (dp, J = 6.5, 3.5 Hz, 2H), 6.93 (dd, J = 8.8, 2.3 Hz, 2H). 13C NMR (101 MHz, DMSO-d6) δ 162.96, 160.81, 146.88, 133.68, 132.93, 132.33, 129.81, 128.49, 128.30, 127.79, 127.03, 126.73, 123.95, 122.74, 115.11. HRMS calculated for C18H15N2O2 (M+H)+ : 291.11280, Found: 291.11284.

### Cell culture

C2C12 cells (ATCC® CRL-1772™), mouse myoblast cell line, were maintained in Dulbecco’s modified Eagle’s medium (DMEM) supplemented with 10% FBS and 1% L-Glutamine. Primary myoblasts were maintained in DMEM:F12 (1:1) media supplemented with 40% heat inactivated FBS and 10% amninomax (Lifetech). Cells were treated with SLU-PP-332 or DMSO (10 µM). After 24 hours of treatment RNA was extracted by Invitrogen Purelink RNA Mini Kit (Invitrogen). All groups were tested in triplicates.

### Co-transfection assays

As previously described^51^, HEK293 cells were maintained in Dulbecco’s modified Eagle’s medium (DMEM) supplemented with 10% fetal bovine serum at 37 °C under 5% CO2. Twenty-four hours prior to transfection, HEK293 cells were plated in 96-well plates at a density of 2 × 104cells/well. GAL4-NR-LBD, or FLAG-ERR-FL plasmids were used in the luciferase assay.

### Real Time PCR (RT-PCR)

The RNA samples were reverse transcribed using the qScript cDNA kit (Quanta). All samples were run in duplicates and the analysis was completed by determining ΔΔCt values. The reference gene used was 36B4, a ribosomal protein gene. Primers sequences are listed on in the supplementary methods.

### Bioenergetic Profile of C2C12 cells

Bioenergetics profile tests in C2C12 myoblasts were conducted as described by Nicholls et al ^52^. The day before (24hrs) the assay, C2C12 cells were seeded (10000/well) in growth media on the 96-well XF Flux Analyzer (SeaHorse®) cell plate.

### Differential Scanning Fluorimetry

ERRγ protein was diluted in a buffer containing 25 mM HEPES pH 7.5, 300 mM NaCl, 10mM DTT, 1mM EDTA at a final concentration of 0.1 mg/mL and mixed with SYPRO-Orange dye (Life technologies S6650). Four different concentrations of ligands (20 µM, 10 µM, 5 µM and 2.5 µM) were used. Six replicate reactions were set up and run in Applied Biosystems Quantstudio 7 Real-Time PCR system. Data were collected at a ramp rate of 0.05°C/s from 24°C through 95°C and analyzed using Protein Thermal Shift Software 1.3.

### Fiber type, SDH and Laminin staining

Fresh cryo-sections (10 μm) were incubated for 1h with Mouse on Mouse (M.O.M, Vector Lab) incubation media and then incubated with BA-D5, SC71 or BF-F3 antibodies (Developmental Studies Hybridoma Bank) for 45 min at 37°C, in PBS-1%BSA. Sections were washed 3 times 5 minutes in PBS, then incubated with secondary antibodies diluted in PBS-1%BSA, for 30 minutes at 37°C. Sections were washed 3 times 5 minutes in PBS and mounted using ProLong Gold mouting media (Thermo Fisher) under glass cover slips.

Fresh cryo-sections (10 μm) were incubated for 30 min at 37C in incubation medium (50mM phosphate buffer, sodium succinate 13.5mg/ml, NBT 10mg/ml in water) placed in a Coplin Jar and then rinsed section in PBS. After staining sections were fixed in 10% formalin-PBS solution for 5 minutes at room temperature and then rinsed in 15% alcohol for 5 minutes. Slides were mounted with an aqueous medium and sealed. Cryostat sections (10 μm) were fixed for 20 minutes in 3% paraformaldehyde in phosphate-buffered saline (PBS), pH 7.4. Sections were blocked with 8% BSA in PBS 1h at RT and then incubated at 4°C overnight with primary antibody for laminin at a 1:200 dilution. Sections were then washed and incubated wit anti-rabbit-FITC (1:1000) for 1 hour at room temperature. Sections were mounted with fluorescent mounting medium containing Dapi (Vector lab) under glass coverslips. All quantifications were performed using ImageJ software.

### Mice

Male C57BL6/J or ob/ob mice were obtained from Jackson Laboratories (Bar Harbor, ME). Studies performed with C57BL6/J or ob/ob mice were approved and conducted in accordance to the Saint Louis University and Washington University Animal Care Use Committees. The conditional ERRα knockout mice used in exercise performance trials have been described^14^. All procedures using the skeletal muscle-specific M-ERRαWT M-ERRα-/-mice were performed in accordance with the City of Hope Institutional Animal Care and Use Committee.

### General mouse studies

For all experiment 8-10 male C57BL6/J mice per group (12 weeks of age for chow) were administered a dose of SLU-PP-332 50 mg/kg (i.p., b.i.d.) or vehicle for 28 days or 12 days. At termination of the experiment tissues were collected for gene expression analysis by real time qPCR using methods previously described. Food intake and body weight were monitored daily in these experiments and body composition was measured prior to initiation and termination of the experiments by NMR (Bruker BioSpinLF50). Plasma was collected for triglyceride and cholesterol measurements. All b.i.d. dosing was performed with dosing occurring at CT0 and CT12.

### Exercise endurance in WT mice

Six male mice (C57BL6/J, 12 weeks old) were either treated with vehicle control (10% tween, 10% DMSO, 80% PBS) or SLU-PP-332 (50mg/kg, i.p.), run on Exer3-/6 treadmill (Columbus Instruments) for 45min at 12m/min or left untreated. Animals were sacrificed by CO2 asphyxiation 1hr, 3hrs or 6hrs after intervention. For exhaustion protocol, 6 males C57BL6/J, 12 weeks old, were with vehicle control (10% tween, 10% DMSO, 80% PBS) or SLU-PP-332 (50mg/kg, IP) for 6 days before testing. Mice were allowed to acclimate to the treadmill for 10 minutes / day, every day at 2m/min. The day of the test, mice were run 1h after the last dose of vehicle or drug. Mice were running for 2 minutes at 10m/min, then 6 minutes at 12m/min then ran until exhaustion by increasing the speed of the belt for 2m/min every 2 minutes^53^. Exhaustion was assessed by mice allowing 10 consecutive 3ms electrical shock without moving. Mice were sacrificed by CO2 asphyxiation just after exhaustion and exhaustion was confirmed by measuring blood glucose.

### Exercise endurance in muscle specific ERRα KO mice

M-ERRαWT and M-ERRα-/-mice (20-24 wk old, 30.7 + .26 g b.w.) were segregated into vehicle-treated or SLU-PP-332-treated groups (n=5/group). Mice were administered (i.p.) vehicle (10% DMSO, 15% Kolliphor® EL in sterile saline) or 25 mg/kg SLU-PP-332 for 15 days. Prior to run performance trial mice were acclimated to the treadmill (Columbus Instruments Exer 3/6 motorized treadmill) for 3 days (10 min at 10 m/min, then 2 min at 15 m/min). To assess aerobic run performance, mice were run at 10, 12.5 and 15 m/min for 3 min at each speed after which the speed was increased 1m/min every 2 min (max speed 28 m/min) until exhaustion. Basal and post-run blood lactate readings were read to confirm exhaustion. Mice were sacrificed and hindlimb muscles collected 24 hr after the run performance test.

### Glucose measurement

Blood was collected by tail snip and glucose was measure when mice reached exhaustuion using OneTouch Ultra®2 glucometer.

### Pharmacokinetic studies

Pharmacokinetic studies of SLU-PP-332 in mice were performed as previously described^54^. Three-month old C57Bl6/J male mice (n=3) were injected (i.p.) at ZT 1 with 30mg/kg of SLU-PP-332 (5% Tween-5% DMSO-90% PBS). Animals were sacrificed by CO_2_ asphyxiation and tissues were collected at 1h, 2h or 4h after administration of the compound (n=4 per time point). Plasma and tissues (liver, quadriceps and brain) were collected and flash frozen and stored at −80°C until analysis. Tissue samples were weighed and placed into Eppendorf tubes. Naïve tissue was used to prepare standard curves in muscle tissue matrix. To each sample or standard tube was added 3-5 stainless steel beads (2-3 mm) and the appropriate volume of cold 3:1 acetonitrile:water (containing 100 ng/mL extraction internal standard SR8278)^55^ to achieve a tissue concentration of 200 mg/mL. Tubes were placed in a bead beater for 2-3 minutes. Samples and standards (100 µL) were plated in a 96-well plate, 150 µL acetonitrile was added to each well and then centrifuged at 3200 rpm for 5 minutes at 4°C. The supernatant (100 µL) was transferred to a 96-well plate, evaporated to dryness under nitrogen, reconstituted with 100 µL of 0.1% v/v formic acid in 9:1 water:acetonitrile, and vortexed for 5 minutes. Plasma samples or standards prepared in plasma matrix (100 µL) were added to a 96-well plate. To each well, 400 µL of cold acetonitrile containing 100 ng/mL extraction internal standard SR8278 was added. The plate was vortexed for 5 minutes at 4oC. The supernatant (300 µL) was transferred to a second 96-well plate, evaporated to dryness under nitrogen, reconstituted with 100 µL of 0.1% v/v formic acid in 9:1 water:acetonitrile, and vortexed for 5 minutes. Finally, to each reconstituted tissue or plasma sample, 10 µL of 1000 ng/mL enalapril in acetonitrile was added as an injection internal standard, and the 96-well plate was vortexed, briefly centrifuged, and submitted for LC/MS analysis. SLU-PP-332 concentrations were determined on a Sciex API-4000 LC/MS system in positive electrospray mode. Analytes were eluted from an Amour C18 reverse phase column (2.1 × 30 mm, 5 µm) using a 0.1% formic acid (aqueous) to 100% acetonitrile gradient mobile phase system at a flow rate of 0.35 mL/min. Peak areas for the mass transition of m/z 291 > 121 for SLU-PP-332, m/z 394 > 189 for the extraction internal standard SR8278, and m/z 376 > 91 for the injection internal standard enalapril were integrated using Analyst 1.5.1 software. Peak area ratios of SLU-PP-332 area/SR8277 area were plotted against concentration with a 1/x-weighted linear regression. Enalapril was used to monitor proper injection signal throughout the course of LC/MS analysis.

### Lipid assays

Plasma triglycerides, total cholesterol and liver enzymes were assessed using an Analox (GM7 MicroStat) instrument and kits provided by the same manufacturer following their protocols.

### Limited proteolysis digestion

In vitro-translated ERRα full length (TNT kit; Promega) was used. Briefly, after incubating at room temperature for 15 min with ligands (1-5-10 µM), receptor proteins were digested at room temperature for 10 min with 10 μg/ml trypsin. The proteolytic fragments were separated on a 4-15% SDS polyacrylamide gel (BioRad) and visualized by Coomassie Blue staining.

### Assessment of locomotor activity

Locomotor activity was assessed using mice housed in cages with free access to running wheels. Briefly, after a 2 days’ acclimation period to wheel-equipped cages in a 12:12 light-dark (LD), locomotor activity was recorded over a 48hrs period. Wheel running data were analyzed using Clocklab software (Actimetrics, Evanston, IL).

### Mitochondrial DNA quantification

Mitochondrial DNA was extracted using QiAamp DNA mini kit following the manufacturer instructions’ (Qiagen). DNA was quantified using Sybr Select Master Mix (Applied Biosystems). All samples were run in duplicates and the analysis was completed by determining ΔΔCt values. The reference gene used was NRDUV1, a genomic DNA marker.

### Statistical Analysis

Data are expressed as mean +/-SEM. Student’s test or Two-way ANOVA were used to calculate statistical significance. P<0.05 was considered significant.

## AUTHOR INFORMATION

### Notes

T.P.B., B.E., and J.K.W. are stockholders in Myonid Therapeutics, Inc., which focuses on ERR based therapeutics.

## Acknowledgments

They also thank Grant Kolar and Barbara Nagel (Saint Louis University) for their help in processing samples for electron microscopy. This work was supported by grants from the NIH (AR069280, MH092769, AG060769 and MH093429; TPB and DK057978, HL105278, HL088093, and ES010337; R.M.E.), and the Leona M. and Harry B. Helmsley Charitable Trust (2017PG-MED001; R.M.E.). R.W. is supported under a T32DK training program, and R.M.E. and M.D. are supported, in part, by a Stand Up to Cancer Dream Team Translational Cancer Research grant and a Program of the Entertainment Industry Foundation (SU2C-AACR-DT-20-16). R.M.E is an investigator of the Howard Hughes Medical Institute and March of Dimes Chair in Molecular and Developmental Biology at the Salk Institute. J.M.H is supported by the American Diabetes Association Innovative Basic Science Award (1-18-IBS-103) and Support from the Lion’s Club Diabetes Innovation Fund.

