## Supplementary figures for "A Synthetic ERRα Agonist Induces an Acute Aerobic Exercise Response and Enhances Exercise Capacity"

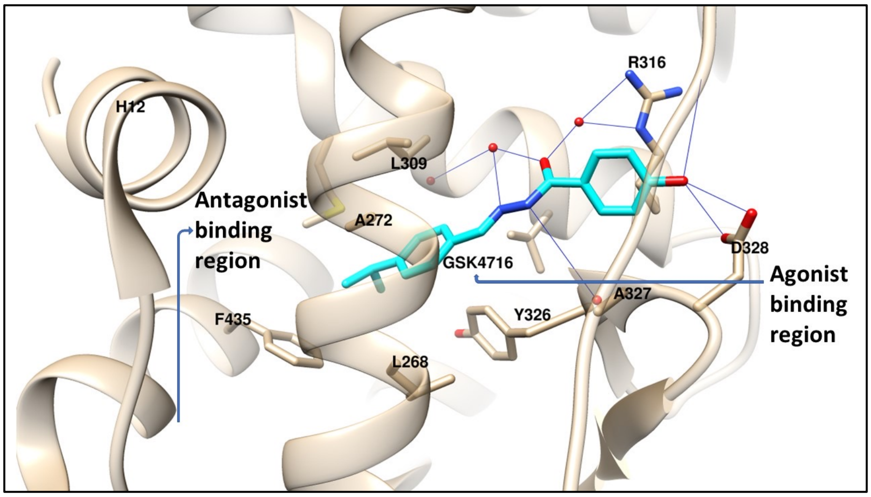


**Supplementary Figure 1.** Ligand interactions in the ERRγ-GSK4716 crystal structure. Hydrogen bonds are shown as blue lines.

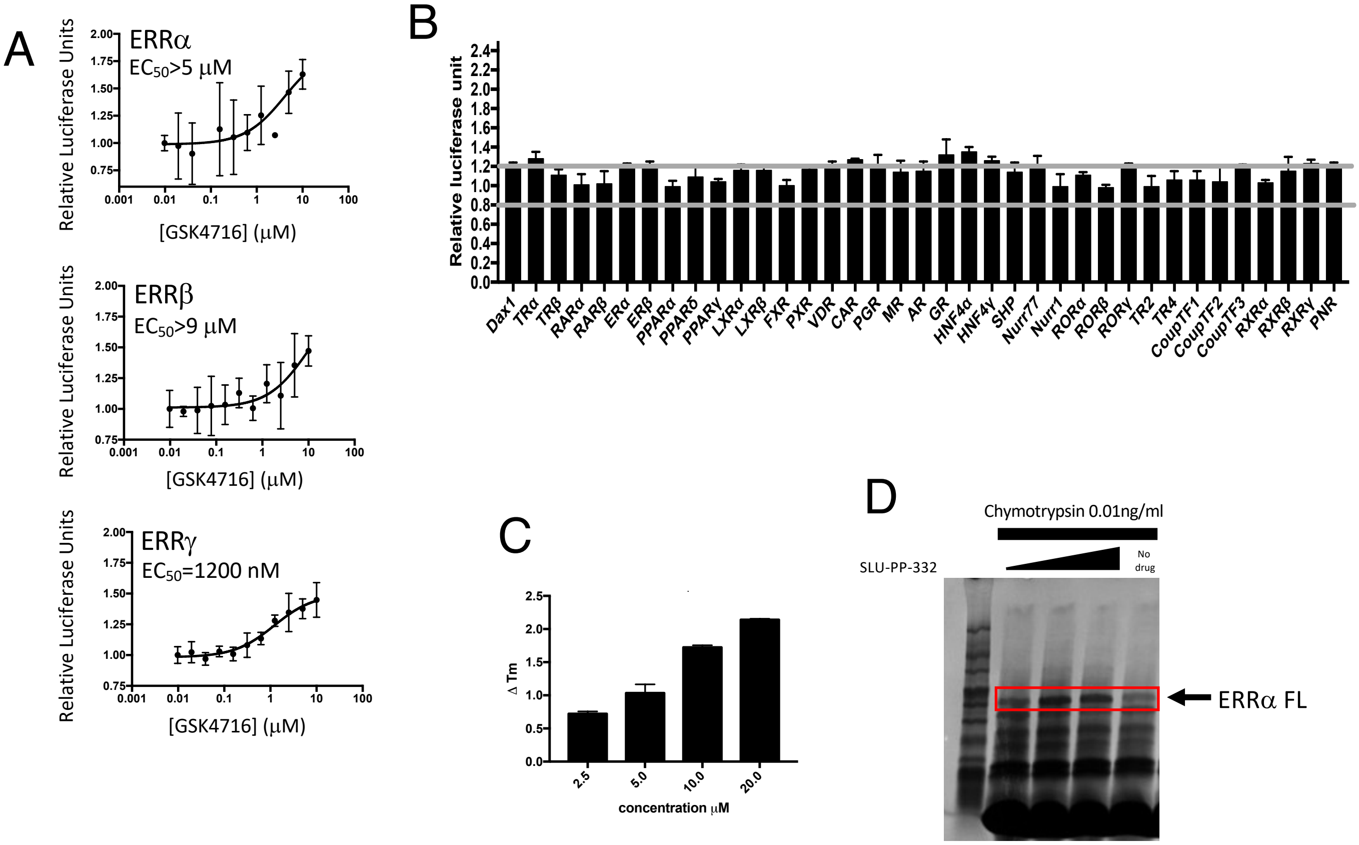


**Supplementary Figure 2. Activity of GSK4716 and Specificity of SLU-PP-332. (A)** Results from a cotransfection assay in HEK293 cells transfected with full length ERRs (α,β,γ) along with an ERRE containing luciferase reporter. **(B)** Specificity assay (in GAL4DBD-LBD receptor format) for a range of nuclear receptors illustrating specificity of SLU-PP-332 (tested at 10 μM) for the ERRs. **(C)** Biochemical assay (thermal shift assay) illustrating that SLU-PP-332 binds directly to the LBD of ERRγ. **(D)** Limited protease digestion assay of ERRα illustrating binding of SLU-PP-332 to the receptor due to protection of the indicated band. Concentrations of SLU-PP-332 were 1, 5 and 10 μM and the full-length ERRα band is indicated. Error bars indicate mean ± s.e.m. and n=3

A

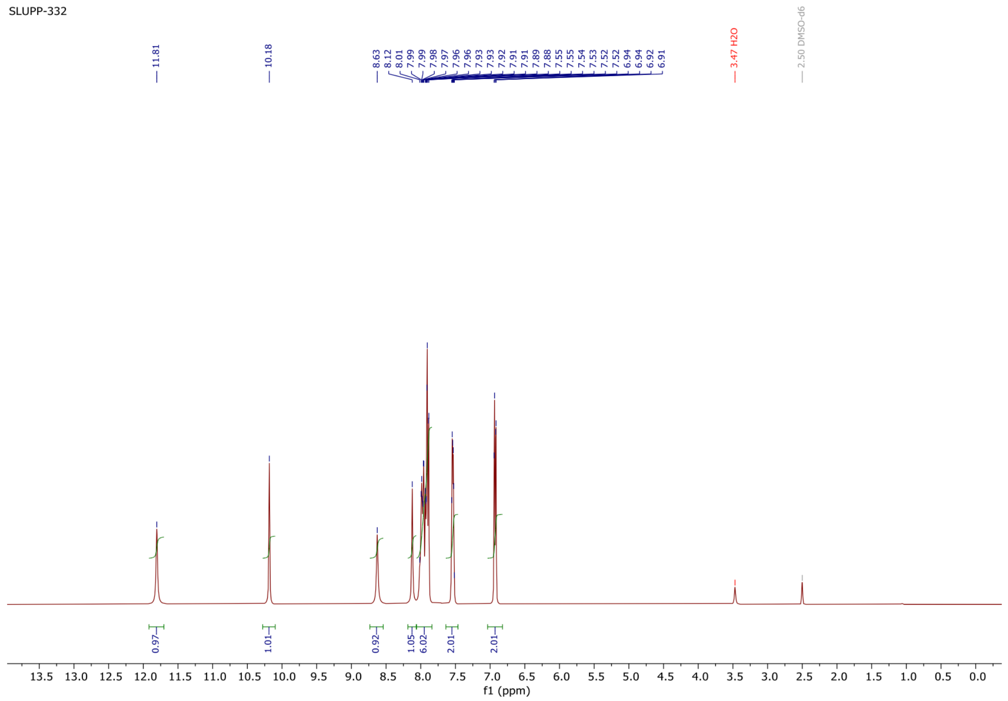


$^{1}H$ NMR Spectrum of SLU-PP-332

B

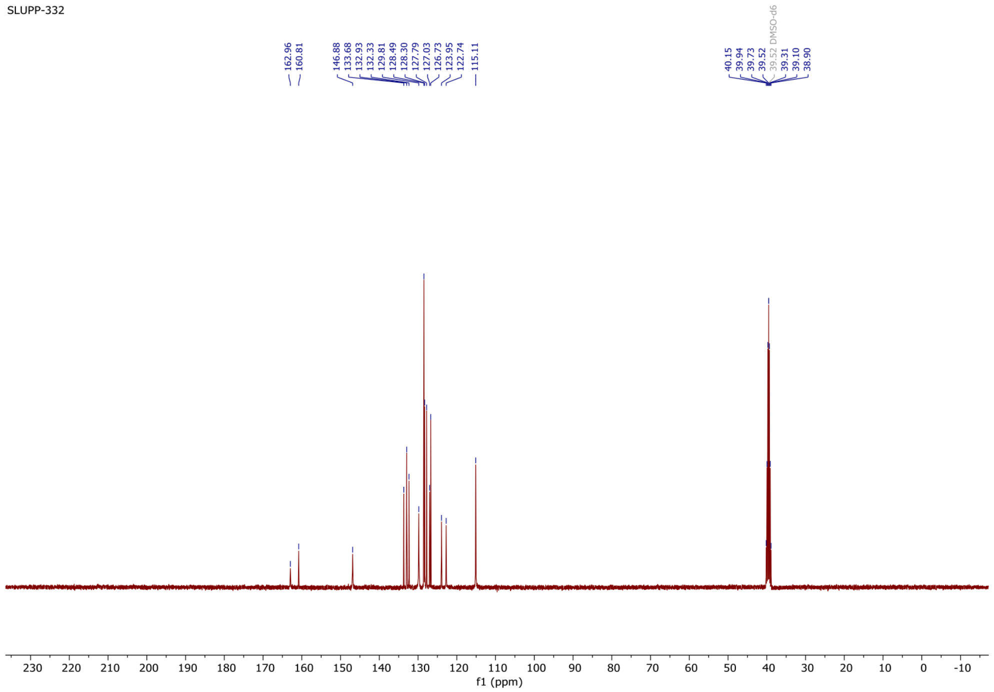


$^{13}C$ NMR Spectrum of SLU-PP-332

C

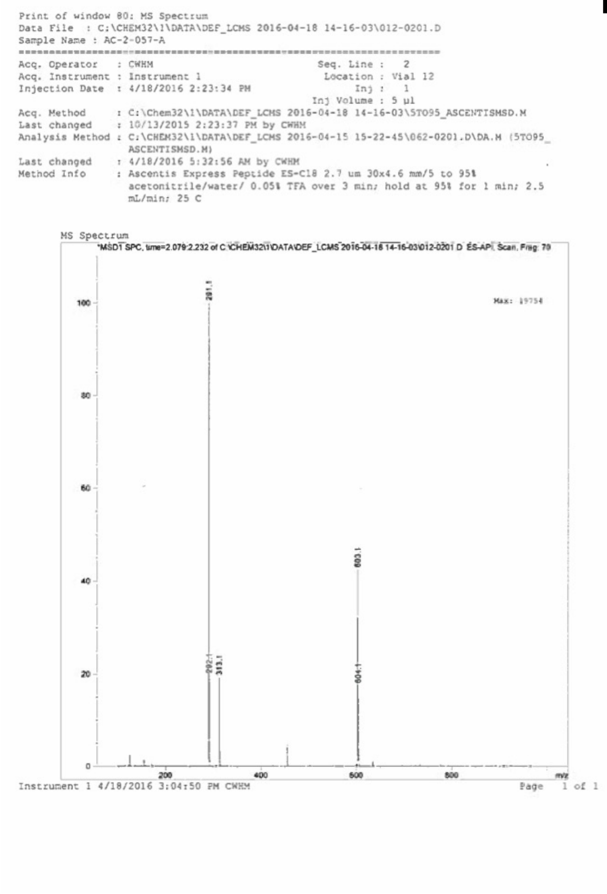


D

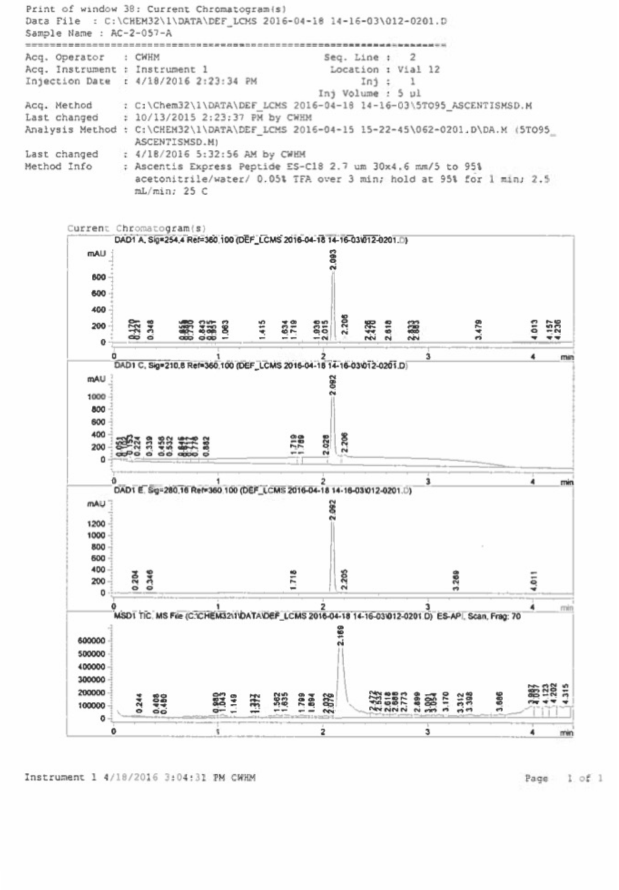


**Supplementary Figure 3. Analytical characterization of SLU-PP-332** A. Sample purity of SLU-PP-332 evidenced by proton NMR **(A)**, 13C NMR **(B)**, mass spectrometry **(C)** and HPLC traces collected at four distinct wavelengths (210, 220, 254 and 280 nM) **(D)**

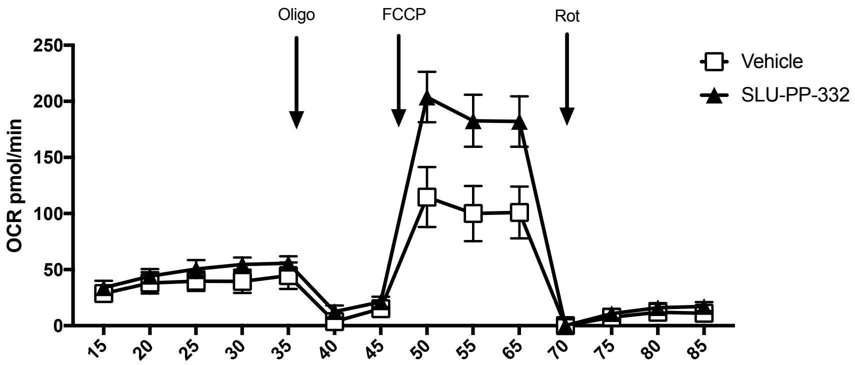


**Supplementary Figure 4. Effects of SLU-PP-332 on mitochondrial respiration in differentiated C2C12 cells.** Analysis of the effect of SLU-PP-332 on mitochondrial respiration in C2C12 cells. Cells were treated for 24h with 10μM of SLU-PP-332 (black triangle) or DMSO (white square) prior the assay.

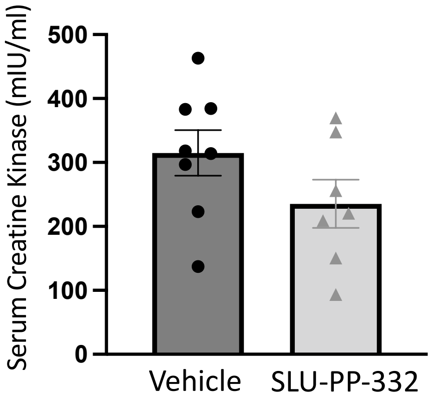


**Supplementary Figure 5. Lack of effect of SLU-PP-332 on serum creatine kinase levels.** As a marker of potential muscle toxicity serum creatine kinase was examined in mice treated chronically (10 days b.i.d.) with 50 mg/kg of SLU-PP-332 i.p.

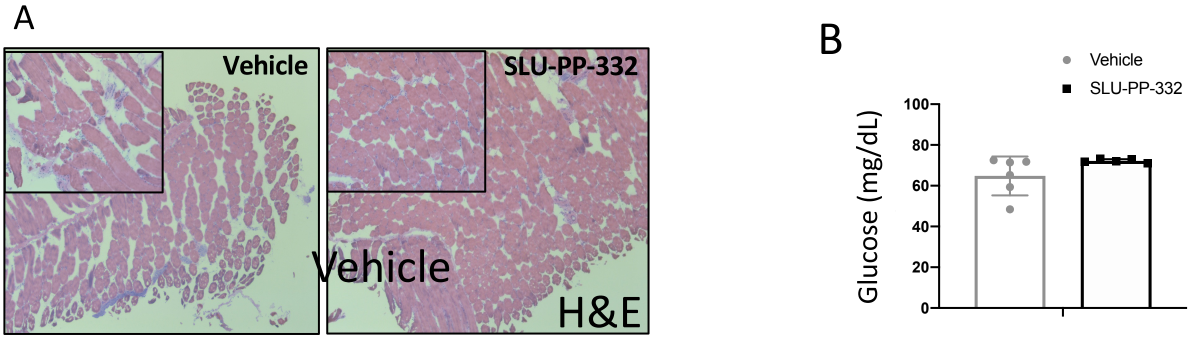


**Supplementary Figure 6. (A)** Histological analysis of hematoxylin and eosin stained skeletal **(B)** Plasma glucose levels from mice exercised to exhaustion either treated with vehicle or SLU-PP-332.

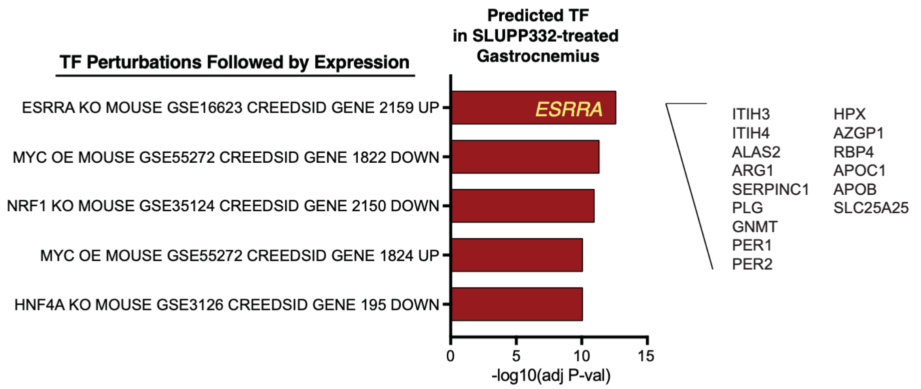


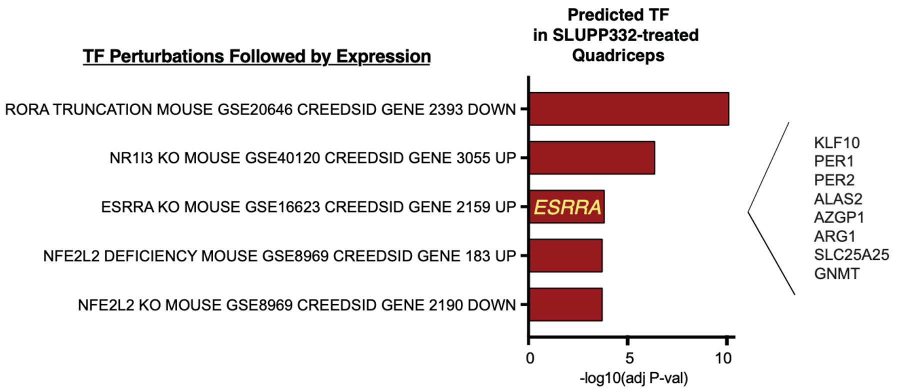


**Supplementary Figure 7. SLU-PP-332 Induces ERRα target genes in skeletal muscle.** Analysis of RNA-seq data obtained from mice treated with SLU-PP-332 reveals regulation of genes that contain ERRα binding sites as revealed us9ing the EnrichR tool.

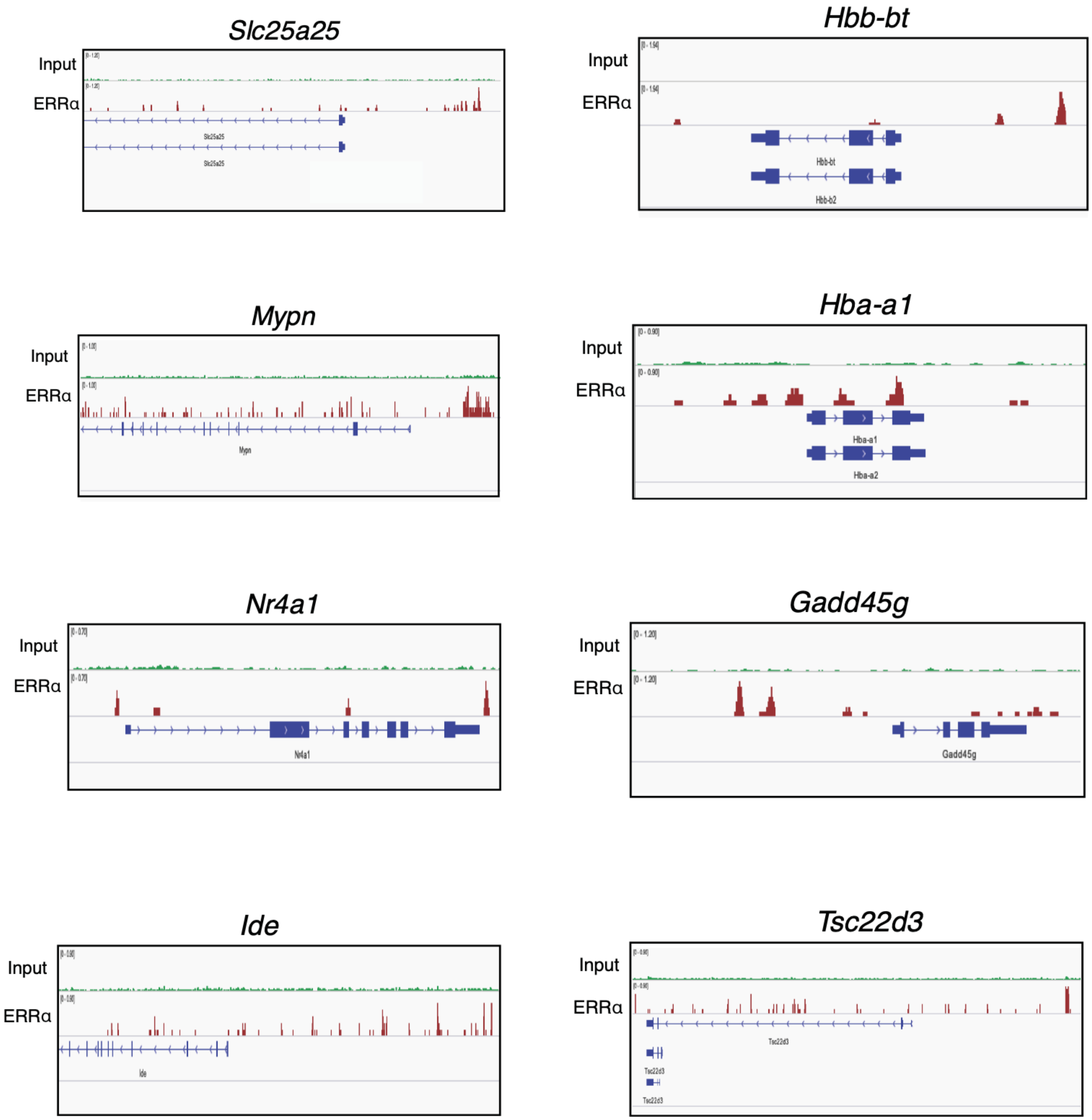


**Supplementary Figure 8**. Genes identified as induced by SLU-PP-332 by RNA seq contain ERRa binding sites. Several of the highest SLU-PP-332 upregulated genes identified in the skeletal muscle of mice contain ERRa binding sites identified in ChIP-seq studies performed in C2C12 cells (Salatino et al., 2016).,

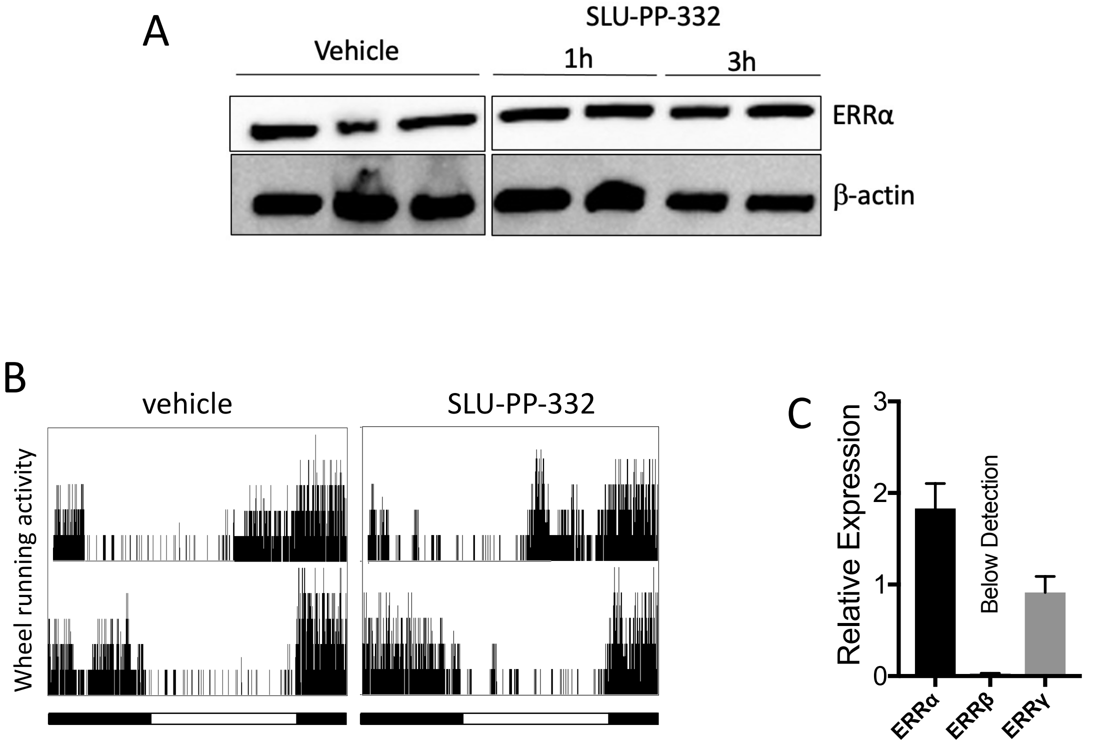


**Supplementary Figure 9. (A)** Western blot illustrating that ERRa protein levels do not change following treatment with SLU-PP-332 (1h or 3h post administration). as described in Fig. 3g **(B)** Locomotor activity of mice treated with SLU-PP-332 reveals no effect on circadian behavior. Light and dark lies represent light and dark periods (12h;12h) **(C)** Relative expression of ERRα, β and γ in primary myocytes.

**Supplementary Table 1**. Contribution of the binding energy components to the total binding free energy ΔG (Kcal/mol).

| Complex | ΔH | TΔS | ΔG = ΔH - TΔS |
|---|---|---|---|
| ERRγ-GSK4716 | -41.4 | -25.3 | -16.1 |
| ERRγ-SLUPP332 | -38.9 | -21.3 | -17.6 |
| ERRα-GSK4716 | -36.9 | -26.8 | -10.1 |
| ERRα-SLUPP332 | -38.3 | -21.6 | -16.7 |

### Supplementary Table 2

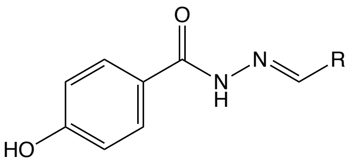


| Compound | R Group | ERRα $EC_{50}$ (μM) |
|---|---|---|
| GSK4716 | 4-isopropyl Phenyl | >1 |
| Cmpd #1 | 4-tertbutyl Phenyl | >1 |
| Cmpd #2 | 4-Cl Phenyl | >1 |
| Cmpd #3 | 1-Napthyl | >1 |
| SLU-PP-332 | 2-Napthyl | 0.098 |
| Cmpd #5 | 6-Hydroxynaphthalen-2-yl | >1 |
| Cmpd #6 | 6-Methoxynaphthalen-2-yl | >1 |

### Supplementary Table 3

#### Clinical Chemistry and CBC of Vehicle and SLU-PP-332 Treated Mice

| | Vehicle | | SLU PP-332 | | C57Bl/6J normal range | |
|---|---|---|---|---|---|---|
| | **Average** | SEM | **Average** | SEM | **Average** | SEM |
| WBC | 6.1775 | 0.9 | 6.84 | 1.4 | 4.5 | 2.5 |
| Neutrophiles | 1038.75 | 152.6 | 1548 | 161.3 | 690 | 390 |
| Lymphocytes | 5101 | 801.5 | 5005.4 | 1315.7 | 3550 | 2000 |
| Monocytes | 34.5 | 12.5 | 267.4 | 68.6 | 200 | 130 |
| Eosinophiles | 1.25 | 1.3 | 3 | 1.0 | 70 | 40 |
| Basophiles | 3 | 2.1 | 18.2 | 11.5 | 10 | 10 |
| RBC | 8.3425 | 0.1 | 7.638 | 0.2 | 8.5 | 0.5 |

| | Vehicle | | SLU PP-332 | | C57Bl/6J normal range | |
|---|---|---|---|---|---|---|
| | **Average** | SEM | **Average** | SEM | **Average** | SEM |
| Hg | 13.4 | 0.1 | 12.3 | 0.3 | 11.6 | 1.0 |
| HT | 45.1 | 0.3 | 40.6 | 0.6 | 36.9 | 2.0 |
| Platelets | 738.0 | 37.4 | 655.8 | 83.7 | 802.0 | 239.0 |

| | Vehicle | | SLU PP-332 | | C57Bl/6J normal range | |
|---|---|---|---|---|---|---|
| mmol/l | **Average** | SEM | **Average** | SEM | **Average** | SEM |
| Sodium | 149.5 | 1.0 | 147.4 | 1.0 | 149.0 | 2.0 |
| Potassium | 7.4 | 0.2 | 7.5 | 0.1 | 4.3 | 0.7 |
| Chhloride | 102.5 | 0.7 | 102.1 | 1.7 | 113.0 | 2.0 |
